# RAxML-NG: A fast, scalable, and user-friendly tool for maximum likelihood phylogenetic inference

**DOI:** 10.1101/447110

**Authors:** Alexey M. Kozlov, Diego Darriba, Tomáš Flouri, Benoit Morel, Alexandros Stamatakis

## Abstract

**Motivation:** Phylogenies are important for fundamental biological research, but also have numerous applications in biotechnology, agriculture, and medicine. Finding the optimal tree under the popular maximum likelihood (ML) criterion is known to be NP-hard. Thus, highly optimized and scalable codes are needed to analyze constantly growing empirical datasets.

**Results:** We present RAxML-NG, a from scratch re-implementation of the established greedy tree search algorithm of RAxML/ExaML. RAxML-NG offers improved accuracy, flexibility, speed, scalability, and usability compared to RAxML/ExaML. On taxon-rich datasets, RAxML-NG typically finds higher-scoring trees than IQTree, an increasingly popular recent tool for ML-based phylogenetic inference (although IQ-Tree shows better stability). Finally, RAxML-NG introduces several new features, such as the detection of terraces in tree space and a the recently introduced transfer bootstrap support metric.

**Availability:** The code is available under GNU GPL at https://github.com/amkozlov/raxml-ng.RAxML-NG web service (maintained by Vital-IT) is available at https://raxml-ng.vital-it.ch/.

## 1 Introduction

RAxML (Stamatakis, 2014) is a popular maximum-likelihood (ML) tree inference tool which has been developed and supported by our group for the last 15 years. More recently, we also released ExaML (Kozlov *et al.*, 2015), a dedicated code for analyzing genome-scale datasets on supercomputers. ExaML implements the core tree search functionality of RAxML and scales to thousands of CPU cores. Other widely-used ML inference tools are, for instance, IQ-Tree (Nguyen *et al.*, 2015), PhyML (Guindon *et al.*, 2010), and FastTree (Price *et al.*, 2010).

Here, we introduce our new code called RAxML-NG (RAxML Next Generation). It combines the strengths and concepts of RAxML *and* ExaML, and offers several additional improvements which we describe in the next section.

## 2 New Features and Optimizations

### Evolutionary model extensions

While RAxML/ExaML only fully supported the General Time Reversible (GTR) model of DNA substitution, RAxML-NG now supports all 22 ‘classical’ GTR-derived models. All model parameters (including branch lengths) can be either optimized or fixed to user-specified values. RAxML-NG also offers the following features: (1) edge-proportional branch length estimation for multi-gene alignments, (2) FreeRate model of rate heterogeneity (Yang, 1995), (3) per-rate scalers in the Γ model of rate heterogeneity to prevent numerical underflow on large trees.

### Search algorithm modifications

The subtree enumeration method used in RAxML/ExaML occasionally skipped promising topological moves; this has now been fixed in RAxML-NG (see Supplementary data for details). Further, RAxML-NG employs a two-step L-BFGS-B method (Fletcher, 1987) to optimize the parameters of the LG4X model (Le *et al.*, 2012). This approach (first introduced in IQ-Tree) is usually faster and more stable than the sequential optimization using Brent’s method in RAxML/ExaML.

### Transfer bootstrap

RAxML-NG can compute the novel branch support metric called transfer bootstrap expectation (TBE) recently proposed in (Lemoine *et al.*, 2018). Compared to the classic Felsenstein bootstrap, TBE is less sensitive to individual misplaced taxa in replicate trees, and thus better suited to reveal well-supported deep splits in large trees with thousands of taxa.

### Phylogenetic terraces

Certain patterns of missing data in multigene alignments can yield multiple tree topologies with identical likelihood scores – a phenomenon known as *terraces* in tree space (Sanderson *et al.*. 2011). RAxML-NG employs the recently released *terraphast* library (Biczok *et al.*, 2017) to assess if the inferred best-scoring ML tree resides on a terrace, and report the size of that terrace.

### Performance and scalability

In RAxML-NG, we further optimized the vectorized likelihood computation kernels and eliminated known sequential bottlenecks of RAxML. We also integrated an optimization technique for likelihood calculations known as *site repeats* (Kobert *et al.*, 2017) which yields runtime improvements of 10% to 60%. Finally, RAxML-NG implements several features for enhancing parallel efficiency, previously only available in ExaML: (1) efficient fine-grained parallelization with MPI or MPI+pthreads, (2) binary input file format (compressed alignment), (3) restart from a checkpoint, (4) improved load balancing for multi-gene alignments (Kobert *et al.*, 2014)

### Usability

Several RAxML-NG features aim to improve usability and avoid common pitfalls: auto-detection of CPU instruction set and number of cores, recommendation for the optimal number of threads, automatic restart from the last checkpoint after program interruption, search progress reporting in the log file etc.

### Modularization

RAxML and ExaML are large monolithic codes. This hindered maintenance, extension, and code reuse. In RAxML-NG, we encapsulated the phylogenetic likelihood kernels and numerical optimization routines in two libraries: *libpll* (https://github.com/xflouris/libpll-2) and *pll-modules* (https://github.com/ddarriba/pll-modules), respectively. Both libraries include unit tests and are also being used by other software tools developed in our lab such as ModelTest-NG and EPA-NG (Barbera *et al.*, 2018). This yields our likelihood computation code more error-proof than in RAxML/ExaML.

## 3 Evaluation

A recent evaluation of fast ML-based methods (Zhou *et al.*, 2018) showed that IQTree yields the best tree inference accuracy, closely followed by RAxML/ExaML. Thus, we benchmarked RAxML-NG against these three programs on the collection of empirical datasets used by Zhou *et al.* RAxML-NG found the best-scoring tree for the highest number of datasets (19/21) among all programs tested, while being 1.3× to 4.5× faster. Furthermore, it scales to the large number of cores with a parallel efficiency of up to 125% (see supplement for details). In summary, RAxML-NG is clearly superior to RAxML/ExaML, and thus we recommend that the users of these codes upgrade as soon as possible. Comparison to IQTree yielded mixed results: although RAxML-NG is generally faster and returns higher-scoring trees on taxon-rich alignments, IQTree results show much lower variance. Hence, on alignments with strong phylogenetic signal, IQTree may require fewer replicate searches than RAxML-NG to find the best-scoring tree.

## 4 Availability and user support

The RAxML-NG source code as well as pre-compiled binaries for Linux and MacOS are available at https://github.com/amkozlov/raxml-ng. RAxML-NG is also available as a web service (maintained by the Vital-IT unit of the Swiss Institute of Bioinformatics) at https://raxml-ng.vital-it.ch/. An up-to-date user manual is available at https://github.com/amkozlov/raxml-ng/wiki. User support is provided via the RAxML Google group at: https://groups.google.com/forum/#!forum/raxml.

## 5 Future Work

In future versions of RAxML-NG, we plan to add site heterogeneity models such as RAxML-CAT (Stamatakis, 2006) and PhyloBayes-CAT (Le *et al.*, 2008), as well as non-reversible context-dependent models of evolution (Baele *et al.*, 2010). Furthermore, we plan to explore orthogonal parallelization schemes (across tree nodes and/or topological moves), for leveraging the capabilities of modern parallel hardware and more efficiently analyzing datasets with thousands of taxa.

## Acknowledgements

We thank Lucas Czech, Pierre Barbera, and members of the RAxML google group for helpful suggestions and testing the beta version of this software. We also thank Fabio Lehmann and Heinz Stockinger for the implementation and support of the RAxML-NG web server. Fast TBE computation code was contributed by Sarah Lutteropp.

