## Supplementary data for "RAxML-NG: A fast, scalable, and user-friendly tool for maximum likelihood phylogenetic inference"

April 23, 2019

#### 1 New features and improvements

In this Section, we briefly describe the most important improvements of RAxML-NG compared to RAxML/ExaML. A detailed, and up-to-date user manual is available at https://github.com/amkozlov/raxml-ng/wiki.

##### 1.1 Evolutionary model extensions

###### 1.1.1 New DNA models with flexible parametrization

For DNA data, RAxML/ExaML fully supports only one model of nucleotide substitution: the general time-reversible (GTR) model. In RAxML, there is also only limited support for the JC69 (Jukes and Cantor, 1969) and K80 (Kimura, 1980) models. Note that, for partitioned analyses, it is not possible to assign different models to distinct partitions (e.g., JC69 to one partition and GTR to another).

In RAxML-NG, we added full support for all 22 'classical' GTR-derived models (Felsenstein, 2004), which can be used in any combinations (i.e., different models can be assigned to distinct partitions). Furthermore, substitution rates as well as equilibrium/stationary frequencies can be fixed to user-specified values, if desired.

###### 1.1.2 Flexible models for multi-state data

Apart from classical DNA, protein, and binary data, RAxML can also analyze arbitrary multi-state sequence data that is commonly used to encode morphological traits. RAxML supported up to 32 such states which had to be encoded via a pre-defined set of characters (0-9 and A-V).

RAxML-NG now extends this functionality by allowing for up to 64 states and a more flexible user-defined encoding. For instance, a 5-state model can be specified by MULTI5_GTR+M{ABCDE}{-}. Here, characters A through E encode the 5 states of the model, and - represents the gap character (unknown state/missing data). Furthermore, it is now also possible to define ambiguous characters for multi-state models to capture sequence uncertainty. For instance, one can use the character X to represent an ambiguous state *A or C*. In this case, the corresponding mapping between characters and models states needs to be defined in an appropriate file:

```
7   5
ABCDEX-
stateA stateB stateC stateD stateE
A   1,0,0,0,0
B   0,1,0,0,0
C   0,0,1,0,0
D   0,0,0,1,0
E   0,0,0,0,1
X   1,0,1,0,0
-   1,1,1,1,1
```

For instance, we recently used this new functionality of RAxML-NG to implement a 55-state model that can account for amino acid side-chain conformation (Perron *et al.*, 2019).

##### 1.1.3 Rate heterogeneity across sites

ExaML only supports the $\Gamma$ model model of rate heterogeneity across sites (RHAS) with a fixed number of 4 rate categories. RAxML additionally supports a proportion of invariant sites model ($I$). Both, RAxML, and ExaML do not allow for specifying distinct RHAS models for different partitions (e.g., *all* partitions have to be assigned either a $\Gamma$ model, or a $I$ model, or a $\Gamma + I$ model, or none).

In addition to $\Gamma$ and $I$, RAxML-NG now also supports the FreeRate RHAS model (Yang, 1995), which does not rely on any a priori rate distribution. The free rate models has been shown to better fit empirical datasets (Kalyaanamoorthy *et al.*, 2017) tha the $\Gamma$ model. Furthermore, RAxML-NG provides full flexibility for partitioned analyses: each partition can be assigned its own, best-fit RHAS model, number of discrete rate categories. Corresponding parameter values (e.g., the $\alpha$ shape parameter of the $\Gamma$ distribution) can also be specified and fixed by the user.

##### 1.1.4 Branch length linkage across partitions

In partitioned analyses, there are three common ways to estimate branch lengths (sometimes called *branch linkage* models):

- **linked**: all partitions share a common set of (global) branch lengths. This is the most simple model with the lowest number of free parameters ($\#branches$). However, it is often considered as being unrealistic, as it is known that, genes (or genomic regions) evolve at different speeds.
- **unlinked**: each partition has its own, independent set of branch lengths. This model allows for the highest flexibility, but it also introduces an extremely large number of additional free parameters ($\#branches * \#partitions$), which yields it prone to overfitting.
- **scaled (proportional)**: a global set of branch lengths is estimated as for the `linked` mode. However, each partition has an individual scaling factor. The per-partition branch lengths are obtained by multiplying the global branch length values with these individual per-partition scalers. This approach represents a compromise that allows to model distinct evolutionary rates across partitions while, at the same time, only moderately increasing the number of free parameters ($\#branches + \#partitions$).

While RAxML/ExaML only supported `linked` and `unlinked` linkage, RAxML-NG now supports all three branch linkage models described above. The linkage model can be selected via the `--brlen` option. A recent simulation study by Duchene *et al.* (2018) showed that the `scaled` branch linkage model offers the best fit across a large number of representative datasets. This confirms the intuition about its 'good' flexibility versus complexity trade-off. Hence, RAxML-NG uses the `scaled` branch linkage model for partitioned analyses by default.

##### 1.1.5 Per-rate $\Gamma$ scalers

In RAxML/ExaML, inferences of extremely large trees with thousands of taxa under the $\Gamma$ model of rate heterogeneity frequently induced numerical underflow (Izquierdo-Carrasco *et al.*, 2011) in the likelihood calculations. In RAxML-NG, this problem is solved by introducing so-called per-rate scalers (see Section 4.2.2 in (Kozlov, 2018) for details). As the usage of per-rate scalers induces a small, yet observable computational overhead (typically 5% to 10% of overall run time), their use is automatically enabled for analyses of large trees comprising $> 2000$ taxa, where this specific numerical underflow problem is likely to occur. The user can override this default behavior with the `rate-scalers` flag.

#### 1.2 Search algorithm modifications

The RAxML-NG tree search heuristic is based on the greedy hill-climbing algorithm previously used in RAxML/ExaML. In short, it iteratively prunes every subtree of the thus far best-scoring tree, and evaluates possible re-insertions of each subtree into neighboring branches, up to a certain maximum distance (re-insertion radius) from the original pruning position (see (Stamatakis, 2006) and Section 4.2.3 in (Kozlov, 2018) for further details). The subtree enumeration procedure employed by RAxML/ExaML can occasionally skip promising tree moves: even if *both* subtrees adjacent to a branch are 'good' (have a better likelihood score than the subtrees assessed so far), only *one* of those two subtrees (the one with the higher score) will be saved for further evaluation (see Figure 1, *left*). In RAxML-NG, we modified the subtree enumeration procedure to ensure that promising moves, that is, *both* subtrees, are never skipped (see Figure 1, *right*).

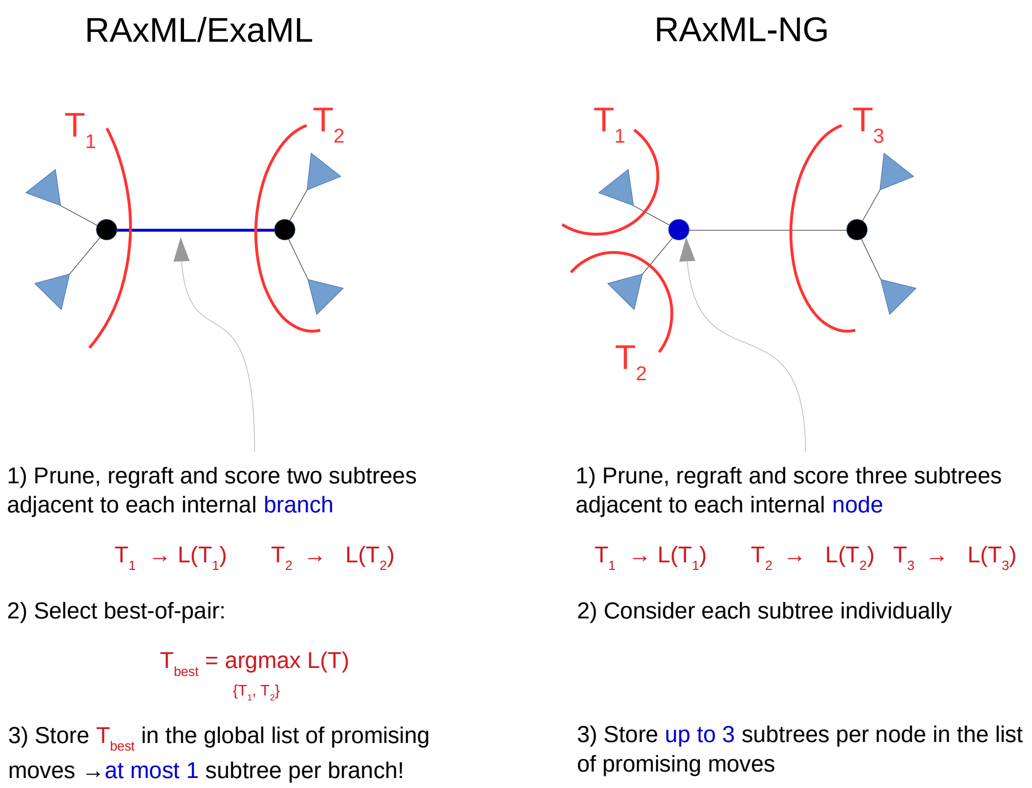


Figure 1: Subtree enumeration strategies in RAxML/ExaML and RAxML-NG.

#### 1.3 Transfer bootstrap

The classical branch support metric used in phylogenetics, *Felsenstein's bootstrap (FBP)*, relies on the strict presence/absence of bipartitions from replicate trees in the best-known ML tree. FBP typically yields low support values for deep branches in extremely large trees with hundreds or thousands of taxa. This is because bipartitions that *exactly* match those in the best-known ML tree are rarely present in the replicates.

The recently introduced *Transfer Bootstrap Expectation (TBE)* metric (Lemoine *et al.*, 2018) attempts to alleviate this problem via a gradual 'transfer' distance. The transfer distance between two branches is defined as the minimum number of taxa that have to be transferred (or removed) to yield the two branches identical (i.e., both branches split the taxon set into identical subsets). The TBE support for a branch in the ML tree is computed based on the *minimum* transfer distance between this branch and *any* branch in a BS replicate tree. In other words, we compare each ML tree branch to its respective closest branch (w.r.t., the transfer distance) in the BS replicate tree (please see Lemoine *et al.* (2018) for further details).

In `RAxML-NG`, we implemented a highly optimized computation of TBE support values. At present, it is $100\times - 500\times$ faster and requires $10\times - 40\times$ less memory than the original implementation in Booster (`https://github.com/evolbioinfo/booster`).

#### 1.4 Phylogenetic terraces

After conducting a tree inference on a partitioned alignment *with unlinked branch lengths*, `RAxML-NG` will automatically check if the resulting tree resides on a phylogenetic terrace (Sanderson *et al.*, 2011). Furthermore, the `--terrace` command allows to check whether a user-specified topology lies on a terrace. Finally, all trees on a terrace can be enumerated and saved in either NEWICK or a compressed NEWICK format (Biczok *et al.*, 2017).

#### 1.5 Performance and scalability

##### 1.5.1 Fine-grained parallelization with MPI and pthreads

RAxML, ExaML, and RAxML-NG all support a fine-grained parallelization of likelihood calculations on a single cadidate tree across alignment sites, that is, alignment sites are distributed across multiple threads/cores. However, in RAxML, this type of parallelization is implemented with pthreads using a fork-join approach, and is thus limited to a single shared-memory node. This precludes the analysis of large concatenated matrices which might require more memory than is available on a single node. This problem was solved in ExaML, which supports a fine-grained parallelization of likelihood calculations across multiple nodes with MPI. Furthermore, ExaML implements an alternative, more efficient parallelization approach which requires fewer synchronizations than the fork-join model. In ExaML, each MPI process runs its own consistent copy of the tree search algorithm, and they only synchronize at two points: 1) tree likelihood evaluation at the virtual root, and 2) likelihood derivative computation during branch length optimization using the Newton-Raphson procedure.

In RAxML-NG, we also implemented this efficient ExaML-style parallelization. It can be used in three configurations: pthreads-only (single shared-memory node), MPI-only, or hybrid MPI+pthreads. Typically, on modern clusters with multicore CPUs, a hybrid configuration with one MPI rank per node and one thread per core yields better parallel efficiency than a pure MPI run with one MPI rank per core.

For 'short' alignments with few sites, a subtatially more coarse-grained parallelization across distinct starting trees or bootstrap replicates could be more efficient. Since RAxML-NG does not natively support such a coarse-grained parallelization (as of version 0.8.1), we recommend using our ParGenes pipeline (Morel *et al.*, 2018) for this purpose which implements an elaborate and efficient resource allocation strategy. Alternatively, one can use custom scripting as shown in the on-line example: https://github.com/amkozlov/raxml-ng/wiki/Parallelization.

##### 1.5.2 Load balancing

In ExaML, we introduced an efficient load balancing algorithm for partitioned alignments (Kobert *et al.*, 2014) which minimizes both, the number of alignment sites, *and* the number of partitions per CPU core. RAxML-NG implements an extended version of this algorithm that includes site weighting support. Site weighting support is necessary to account for partitions with distinct rate heterogeneity models and therefore a varying amount of computations per site. For instance, using a GTR+G4 model on one partition requires approximately 4× more memory and floating point operations than a plain GTR model on another partition. Hence weighting is required to avoid suboptimal parallel load balance.

##### 1.5.3 Checkpointing

Yet another feature that was available in ExaML but not in RAxML is *checkpointing*, that is, the ability to resume a tree search following termination prior to completion for whatever reason, from the stage at which the program was interrupted. This ability is particularly important for long-running analyses in cluster environments where jobs have limited execution time slots (typically, 24 to 48 hours). Compared to ExaML, checkpointing in RAxML-NG is easier to use: it is sufficient to re-run the program without any command line modifications as it will automatically find the latest checkpoint, if present.

##### 1.5.4 Hardware autodetection and resource estimation

In RAxML, a separate binary had to be compiled for every combination of vector instruction set (e.g., AVX) and parallelization type (e.g. pthreads). Albeit the compilation process is described in the RAxML manual, selecting the optimal version was still challenging for users without a strong computational background. Hence, in RAxML-NG we radically simplified the user interface and now only provide a single executable which supports all three parallelization types (pthreads, MPI, and hybid MPI+pthreads) and can also auto-detect the most advanced vector instruction set supported by the CPU it is running on. Furthermore, RAxML-NG can automatically estimate the required amount of memory and the recommended number of threads for the dataset at hand (using --parse command).

### 2 Evaluation

#### 2.1 Experimental setup

##### 2.1.1 Test system configuration

We executed all benchmarking runs on our institutional cluster at the Heidelberg Institute for Theoretical Studies. It consists of 224 compute nodes with dual-slot Intel Haswell CPUs (see Table 1). We compared `RAxML-NG` to three competing state-of-the-art ML tree inference tools: `IQTree`, `RAxML`, and `ExaML` (see Table 2).

| System | HITS cluster | | |
|---|---|---|---|
| Hardware | | Software | |
| CPU model | 2 × Xeon E5-2630 v3 | OS | CentOS Linux release 7.2.1511 |
| CPU architecture | Haswell | | |
| Cores | 16 @ 2.40 GHz | Compiler | GCC 6.4.0 |
| Memory size | 64GB DDR4 | MPI | Open MPI 2.1.1 |

Table 1: Hardware and software specifications of the test system used for benchmarking.

| Tool | Version | Release date | References |
|---|---|---|---|
| `ExaML` | 3.0.19 | May 2017 | (Kozlov *et al.*, 2015; Stamatakis and Aberer, 2013) |
| `IQTree` | 1.6.7 | August 2018 | (Chernomor *et al.*, 2016; Nguyen *et al.*, 2015) |
| `RAxML` | 8.2.10 | March 2017 | (Stamatakis, 2014) |
| `RAxML-NG` | 0.6.0 | June 2018 | (Kozlov, 2018) |

Table 2: ML inference tools including vesion numbers and release dates used for benchmarking.

##### 2.1.2 Datasets

For evaluating `RAxML-NG`, we selected 22 empirical protein and DNA datasets with varying number of taxa (36 up to 1,879), alignment length (18,328 up to 37,350,521 sites), and partition count (1 to 4,116, see Table 3). In particular, we re-analyzed 19 datasets from a recent ML inference tool benchmarking study by Zhou *et al.* (2018) as well as three additional taxon-rich alignments from (Gitzendanner *et al.*, 2018; Shi and Rabosky, 2015; Stamatakis *et al.*, 2010).

For all datasets, we used the original partitioning scheme and substitution models as provided and used in the respective studies.

All multiple sequence alignments used in this study as well as analysis results are available under `https://figshare.com/s/6123932e0a43280095ef`

| Designator | Data type | # taxa | # alignment sites | # unique patterns | # partitions | Reference |
|---|---|---|---|---|---|---|
| SongD1 | DNA | 37 | 1,338,678 | 746,408 | 1 | (Song *et al.*, 2012) |
| MisoD2a | DNA | 144 | 1,240,377 | 1,142,662 | 100 | (Misof *et al.*, 2014) |
| MisoD2b | DNA | 144 | 413,459 | 371,434 | 50 | (Misof *et al.*, 2014) |
| WickD3a | DNA | 103 | 436,077 | 422,676 | 14 | (Wickett *et al.*, 2014) |
| WickD3b | DNA | 103 | 290,718 | 277,375 | 8 | (Wickett *et al.*, 2014) |
| XiD4 | DNA | 46 | 239,763 | 165,781 | 1 | (Xi *et al.*, 2014) |
| PrumD6 | DNA | 200 | 394,684 | 236,674 | 75 | (Prum *et al.*, 2015) |
| TarvD7 | DNA | 36 | 21,410,970 | 8,520,738 | 1 | (Tarver *et al.*, 2016) |
| PeteD8 | DNA | 174 | 3,011,099 | 2,248,590 | 4,116 | (Peters *et al.*, 2017) |
| ShiD9 | DNA | 815 | 20,364 | 13,311 | 29 | (Shi and Rabosky, 2015) |
| StamD10 | DNA | 436 | 1,371 | 1,011 | 1 | (Stamatakis *et al.*, 2010) |
| NagyA1 | AA | 60 | 172,073 | 156,312 | 594 | (Nagy *et al.*, 2014) |
| MisoA2 | AA | 144 | 413,459 | 406,963 | 479 | (Misof *et al.*, 2014) |
| WickA3 | AA | 103 | 145,359 | 144,342 | 11 | (Wickett *et al.*, 2014) |
| ChenA4 | AA | 58 | 1,806,035 | 1,547,914 | 1 | (Chen *et al.*, 2015) |
| StruA5 | AA | 100 | 189,193 | 178,600 | 1 | (Struck *et al.*, 2015) |
| BoroA6 | AA | 36 | 384,981 | 376,803 | 831 | (Borowiec *et al.*, 2015) |
| WhelA7 | AA | 70 | 59,725 | 58,419 | 210 | (Whelan *et al.*, 2015) |
| YangA8 | AA | 95 | 504,850 | 476,259 | 1,122 | (Yang *et al.*, 2015) |
| ShenA9 | AA | 96 | 609,899 | 573,199 | 1 | (Shen *et al.*, 2016) |
| KatzA10 | AA | 798 | 34,991 | 34,937 | 1 | (Katz and Grant, 2014) |
| GitzA12 | AA | 1,897 | 18,328 | 18,303 | 1 | (Gitzendanner *et al.*, 2018) |

Table 3: Characteristics of the datasets used for evaluating `RAxML-NG`.

##### 2.1.3 Evaluation strategy

For each dataset and ML inference tool, we conducted 10 independent searches for the best-known ML tree either using distinct random number seeds (for `IQTree`) or distinct starting trees (for `RAxML`, `ExaML`, and `RAxML-NG`; here we used 5 random and 5 parsimony-based starting trees). Thereby we obtained 40 ML trees for each dataset (10 per inference tool). Subsequently, we re-calculated the ML score (after re-optimizing branch lengths and model parameters on fixed tree topologies) of all final trees with `IQTree` to ensure that all the ML scores of all 40 trees are comparable. Note that, ML values typically differ among distinct inference tools due to different round-off error propagation or subtle differences in the numerical implementation of model parameter optimization routines. Thus, in order to avoid a possible bias toward ML scores obtained by our ML inference tools, we chose to re-evaluate all final trees with `IQTree`.

The ML tree with the (re-evaluated) highest log-likelihood score for a particular dataset is denoted as the *best-known (tree) topology* for that dataset. We assess the relative tree search efficiency of the four ML inference tools by comparing the ML scores of their respective final trees against this best-known ML tree (see Section 2.2.1).

The command lines used for the tree searches and the ML score re-evaluation are provided in Table 4.

Due to technical limitations, we were unable to evaluate some dataset/inference tool combinations. In particular, on the larger benchmark datasets the RAxML searches failed to converge within the cluster job time limit of 24 hours. Given the lack of checkpointing capabilities in RAxML, this prevented us from completing the respective tree inferences. Further, we were not able to analyze the StruA5 dataset with ExaML, since ExaML does not support models with a proportion of invariant sites. This model, more specifically, $LG + \Gamma + I$, was used in the original study by Struck *et al.* (2015). As RAxML and ExaML showed highly similar search efficiencies on the remaining datasets, excluding few aforementioned data points does therefore not affect the overall results.

| Mode | Tool | Command line |
|---|---|---|
| Search | IQTree | iqtree -s <MSA> -q <PARTITIONS> -seed <RSEED> -nt 16 -pre <OUTDIR> |
| Search | RAxML | raxmlHPC-PTHREADS-AVX2 -T 16 -m GTRGAMMA -p <RSEED> -n <RUNNAME> -s <MSA> -q <PARTITIONS> -t <START_TREE> -w <OUTDIR> |
| Search | ExaML | examl-AVX -m GTRGAMMA -n <RUNNAME> -s <BINARY_MSA> -t <START_TREE> -w <OUTDIR> |
| Search | RAxML-NG | raxml-ng –search –msa <MSA> –model <PARTITIONS> –tree <START_TREE> –seed <RSEED> –prefix <OUTDIR> –threads 16 –site-repeats on |
| Evaluation | IQTree | iqtree -s <MSA> -q <PARTITIONS> -te <ML_TREE> -seed <RSEED> -nt 16 -pre <OUTDIR> |

Table 4: Specific command lines used for tree search and log-likelihood evaluation.

#### 2.2 Results

##### 2.2.1 Search Efficiency

Table 5, Figure 3, and Figure 2 show the relative tree search performance of `RAxML-NG` in comparison to other state-of-the-art programs (`RAxML`, `ExaML`, and `IQTree`).

For the MisoD2a dataset, we were unable to reliably determine the best-known tree using our tree scoring procedure (see Section 2.1.3). More specifically, model parameter and branch length re-optimization with `IQTree` on the respective final ML topologies yielded substantially worse log-likelihood scores than those that were originally reported at the end of the tree searches by the tree inference tools. Most importantly, this inconsistency in log-likelihood scores after re-optimization altered the tree ranking. This even holds for ML trees inferred with `IQTree` itself. Interestingly, this issue is not specific to `IQTree` optimization procedure, since we observed similar behavior with `ExaML` and `RAxML-NG` as well. This phenomenon warrants further in-deep investigation, but for the sake of this paper, we simply consider the MisoD2a dataset to be inconclusive and exclude it from our search efficiency analysis.

For 10 out of 21 conclusive datasets, all tested tree inference tools find the best-known ML tree in *at least one* tree search. Moreover, for five out of those 10 'easy' datasets (SongD1, XiD4, BoroA6, WhelA7, and YangA8), the phylogenetic signal was so strong that the best-known ML tree was recovered in *every* single tree search by *any* tool.

The remaining datasets are more challenging and thus allow to assess the performance differences between the tools. None of the tools was able to find best-known tree for all 21 conclusive datasets. `RAxML-NG` showed the best result (19/21), followed by `IQTree` (17/21). In general, `RAxML-NG` (and, to a lesser extent, `RAxML`/`ExaML`) tends to outperform `IQTree` on taxon-rich datasets (ShiD9, KatzA10, and GitzA12). This may indicate that the lazy SPR moves employed by `RAxML-NG`, `RAxML`, and `ExaML` might be better suited to explore larger tree search spaces than the more local NNI moves of `IQTree`, despite the randomized perturbations steps used in `IQTree` for escaping local NNI optima. On the other hand, `IQTree` shows better result stability: it found the best-known ML tree in *all* 10 tree searches for 14 datasets (`RAxML-NG`: 7). This could be attributed to the fact that `IQTree` uses multiple starting trees and maintains a collection of promising trees during each individual search, which allows it to more efficiently navigate out of local optima.

In order to reduce the probability of getting stuck in a local optimum, `RAxML-NG` v0.8.0 and later will conduct 20 independent searches by default, using 10 random and 10 parsimony-based starting trees. Despite substantially longer runtimes, this strategy is much more reliable than a single tree search for most datasets (see Figure 3 and Figure 2).

##### 2.2.2 Inference Times

Absolute tree inference times for all datasets and tools are shown in Figure 4. Furthermore, we provide the relative speedup of `RAxML-NG` compared to other tools in Table 6.

`RAxML-NG` was the fastest tool on 20 out of 22 datasets, with speedups ranging from 1.07 (vs. `IQTree` on GitzA12) to 4.53 (vs. `RAxML` on WhelA7). On the remaining 2 datasets, `RAxML-NG` was slower than `ExaML` and/or `RAxML`, but found trees with better ML scores (see Figure 2, Figure 3, and Table 5). Conversely, on the datasets where all four tools found the best-scoring tree in every replicate search, `RAxML-NG` was consistently faster: $1.74\times$–$4.53\times$ compared to `RAxML`, $1.36\times$–$2.00\times$ compared to `ExaML`, and $2.38\times$–$3.20\times$ compared to `IQTree`.

`RAxML-NG` is able to outperform the competing tools because it relies on highly optimized phylogenetic likelihood kernels (implemented in *libpll*), as well as efficient parallelization and load balancing techniques (see main text).

Finally, we should note that although `IQTree` is consistently slower than `RAxML-NG` for an individual tree search, this is somewhat compensated by its higher stability. In other words, a single tree search with `IQTree` has a higher probability of finding the best-scoring tree compared to `RAxML-NG`. Still, on taxon-rich datasets (ShiD9, StamD10, KatzA10 and GitzA12) even `IQTree` shows substantial variation in inference results. Therefore, relying on the results from a single tree search is a risky strategy for all tools evaluated in this study.

##### 2.2.3 Scalability

On large partitioned multi-gene alignments, `RAxML-NG` scales almost linearly up to 1,024 cores (see Figure 5). On DNA datasets (TarvD7 and PeteD8), we measured superlinear speedups of $> 120\%$ in the optimal parallelization regime ($5,000-8,000$ DNA alignment patterns per core). On protein datasets, this effect is less pronounced, but there is still a noticeable increase of parallel efficiency when $1,500 - 2,000$ AA alignment patterns are assigned to each core.

It is worth noting that, the parallel efficiency of `RAxML-NG` is similar (and partially superior) to `ExaML`, our previous dedicated tree inference tool for large clusters and supercomputers. This is noteworthy as the absolute runtimes of `RAxML-NG` are $\approx 1.5\times$ lower and hence, for instance I/O, startup, and communication costs contribute more to overall run-time according to Amdahl's law.

Finally, we assess the scalability of the MPI-enabled version of `IQTree` on the same four large datasets. Please note that the `IQTree-MPI` parallelization strategy (across tree moves) is conceptually different from the parallelization strategies implemented in `ExaML`/`RAxML-NG` (across alignment sites). For this reason, a direct scalability comparison between the tools is difficult. Furthermore, even if `IQTree-MPI` is run with identical random number seeds, using different numbers of cores *can* yield distinct final ML trees. Thus, as the results of parallel runs with different core counts may defer, it is unclear how to quantify parallel efficiency.

| Dataset | ML trees searches which found the best-known tree | | | |
|---|---|---|---|---|
| | `IQTree` | `RAxML` | `ExaML` | `RAxML-NG` |
| SongD1 | 10 | 10 | 10 | 10 |
| MisoD2a* | NA | NA | NA | NA |
| MisoD2b | 10 | 0 | 0 | 5 |
| WickD3a | 10 | 0 | 0 | 10 |
| WickD3b | 10 | 0 | 0 | 9 |
| XiD4 | 10 | 10 | 10 | 10 |
| PrumD6 | 10 | 0 | 0 | 6 |
| TarvD7 | 10 | 8 | 8 | 9 |
| PeteD8 | 0 | 0 | 0 | 4 |
| ShiD9 | 0 | 0 | 0 | 1 |
| StamD10 | 1 | 0 | 0 | 0 |
| NagyA1 | 10 | 4 | 4 | 5 |
| MisoA2 | 10 | NA | 0 | 5 |
| WickA3 | 10 | 8 | 8 | 4 |
| ChenA4 | 10 | 7 | 7 | 10 |
| StruA5 | 2 | 0 | NA | 9 |
| BoroA6 | 10 | 10 | 10 | 10 |
| WhelA7 | 10 | 10 | 10 | 10 |
| YangA8 | 10 | 8** | 10 | 10 |
| ShenA9 | 10 | 9 | 9 | 8 |
| KatzA10 | 0 | NA | 1 | 0 |
| GitzA12 | 0 | NA | 0 | 1 |
| | Datasets for which the best-known tree was found | | | |
| | 17 | 10 | 11 | 19 |

Table 5: Number of ML tree searches (out of 10) which yielded the best-known tree per dataset and inference tool. * Best-known tree cannot be determined for this dataset (see explanation in text). ** Two `RAxML` tree searches exceeded the cluster job runtime limit.

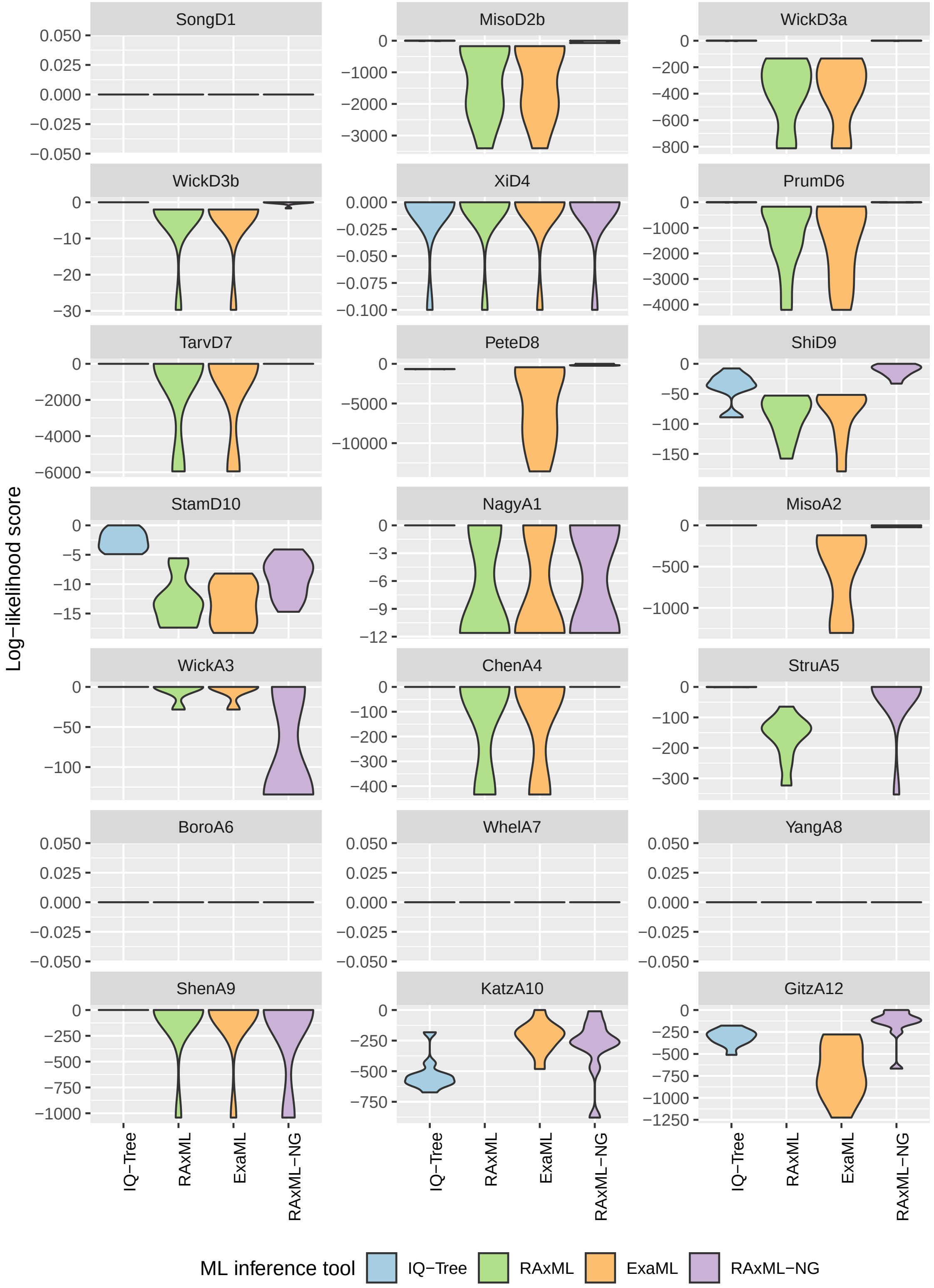


Figure 2: Log-likelihood difference to the best-known tree (normalized log-likelihood score).

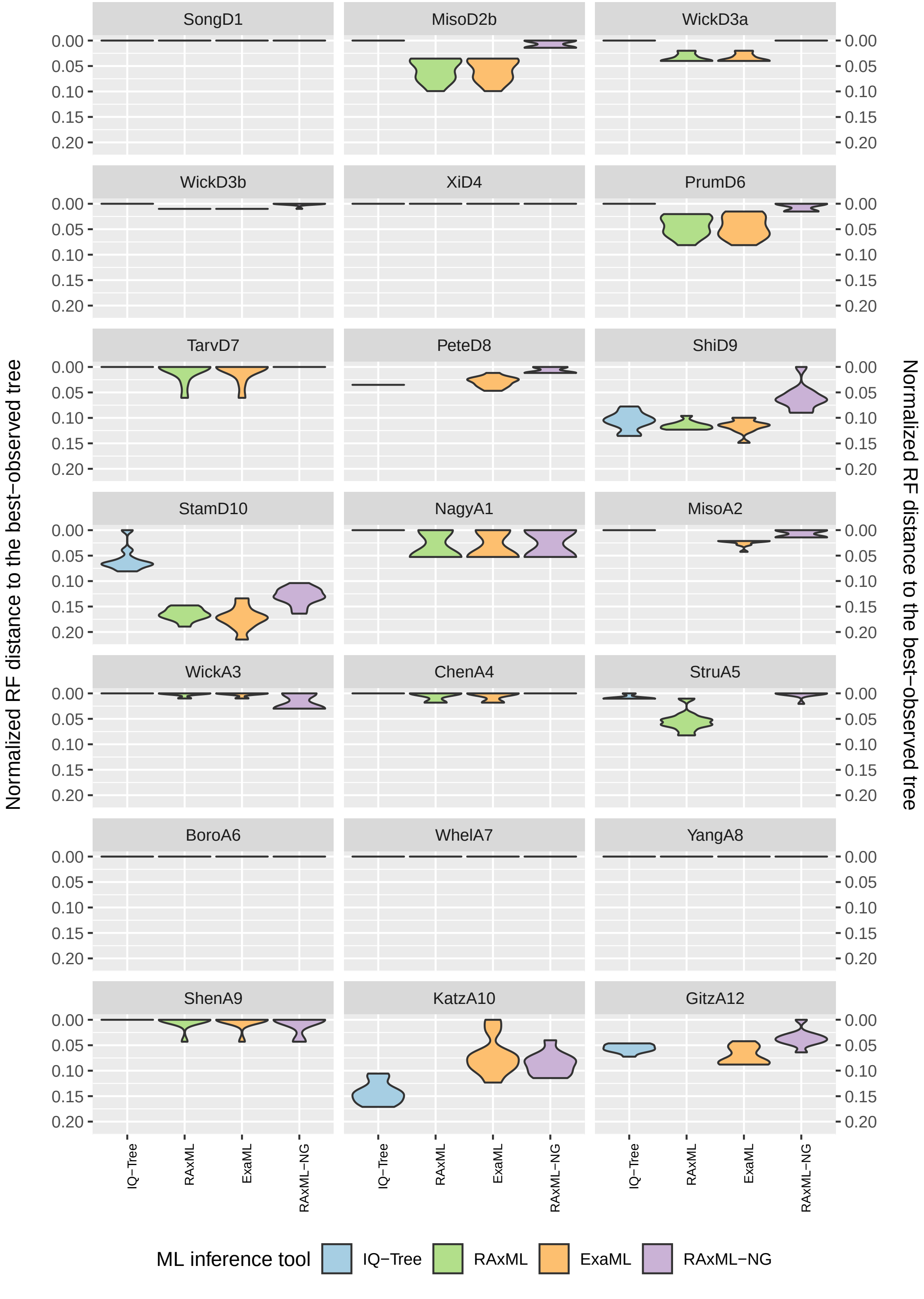


Figure 3: Normalized Robinson-Foulds (RF) distance to the best-known ML tree.

| Dataset | RAxML-NG speedup (x) compared to | | |
|---|---|---|---|
| | RAxML | ExaML | IQTree |
| SongD1 | 1.79 | 1.37 | 3.20 |
| MisoD2a | NA | 2.16 | 2.47 |
| MisoD2b | 2.77 | 1.73 | 4.04 |
| WickD3a | 1.68 | 1.25 | 2.59 |
| WickD3b | 1.82 | 1.33 | 1.64 |
| XiD4 | 1.74 | 1.46 | 2.72 |
| PrumD6 | 1.86 | 1.24 | 2.53 |
| TarvD7 | 1.86 | 1.57 | 3.55 |
| PeteD8 | NA | 1.14 | 1.95 |
| ShiD9 | 1.47 | 0.94 | 3.86 |
| StamD10 | 0.91 | 0.57 | 6.58 |
| NagyA1 | 3.73 | 1.65 | 1.53 |
| MisoA2 | NA | 1.93 | 1.54 |
| WickA3 | 1.66 | 1.43 | 2.11 |
| ChenA4 | 2.11 | 1.40 | 2.18 |
| StruA5 | 3.11 | NA | 1.09 |
| BoroA6 | 2.36 | 1.36 | 2.88 |
| WhelA7 | 4.53 | 2.00 | 2.50 |
| YangA8 | 2.95 | 1.86 | 2.38 |
| ShenA9 | 1.78 | 1.32 | 1.18 |
| KatzA10 | NA | 1.71 | 1.12 |
| GitzA12 | NA | 1.09 | 1.07 |

Table 6: Average RAxML-NG speedups relative to RAxML, ExaML, and IQTree. NA = this dataset cannot be analyzed due to job runtime limit (RAxML) or lack of Γ+I model support (ExaML).

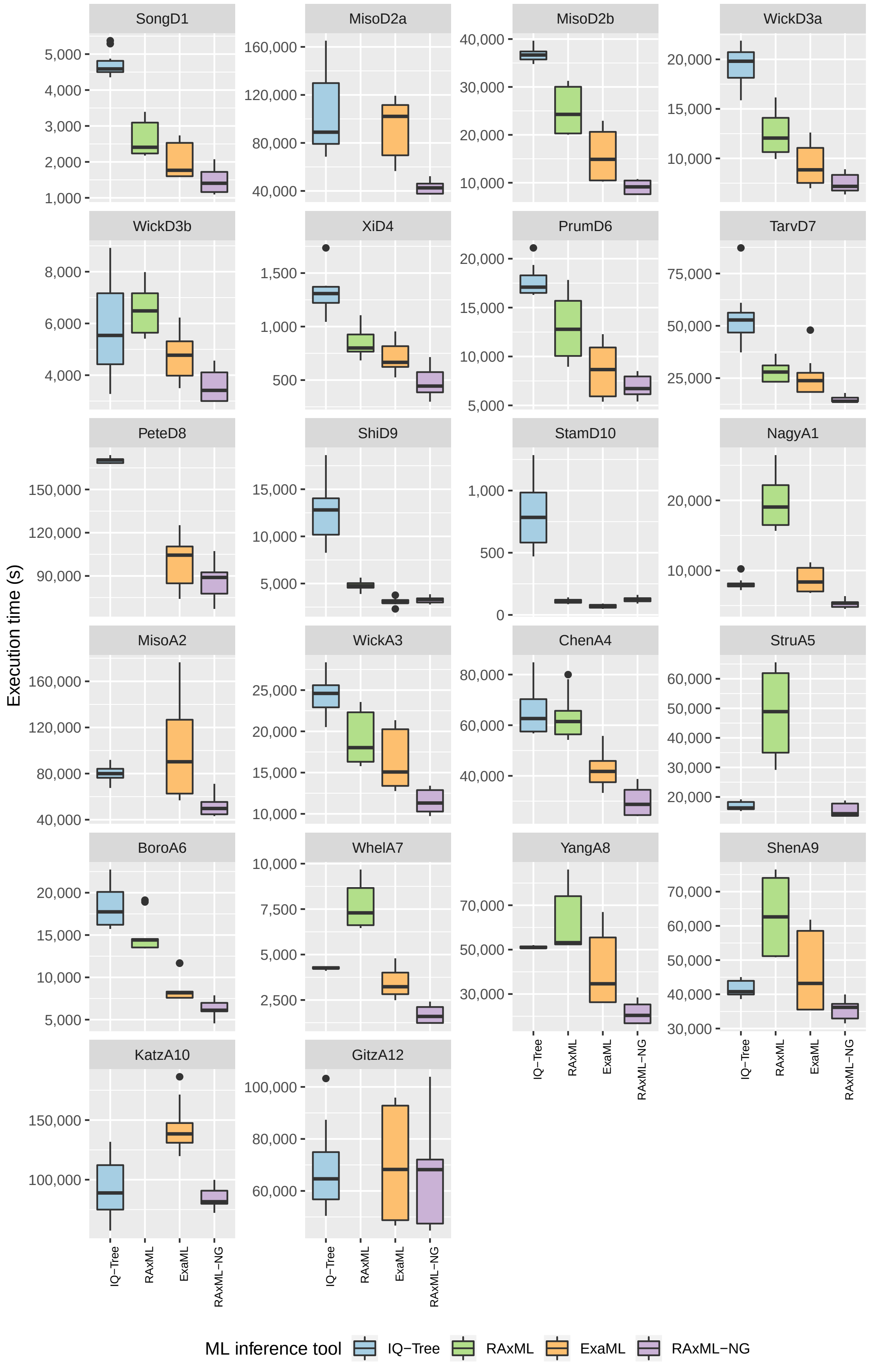


Figure 4: Wall-clock execution times in seconds (16 threads / 1 compute node).

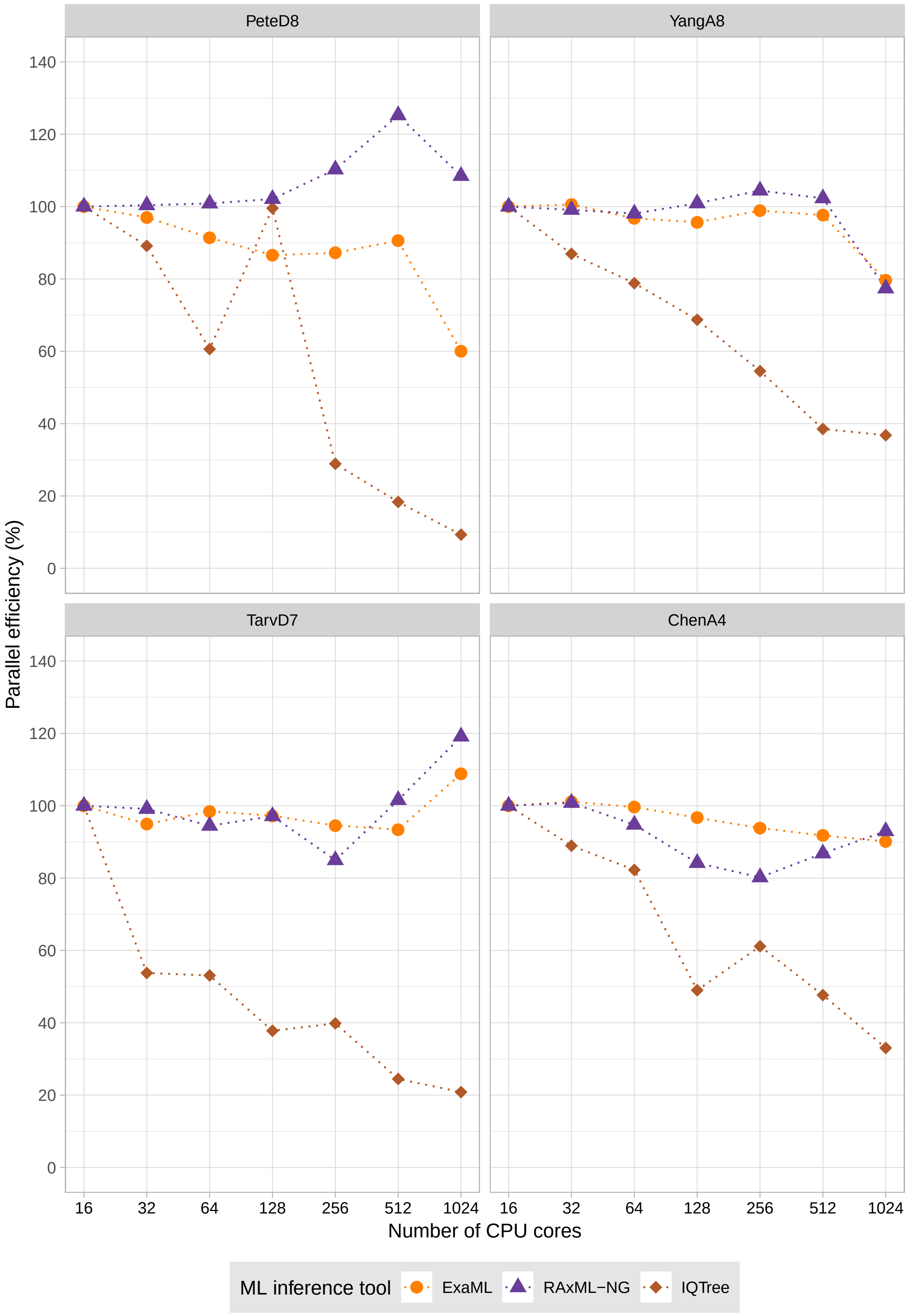


Figure 5: Strong scaling efficiency of `RAxML-NG` vs. `ExaML` on large phylogenomic datasets. On PeteD8 dataset, `IQTree-MPI` runs converged to distinct final ML trees.

.
